# Microstructure of white matter association tracts in the temporal lobe predicts the timing of electrophysiological activity and behaviour in picture naming

**DOI:** 10.64898/2026.10.26.761817

**Authors:** Irina Chupina, Natascha M. Roos, Joanna Sierpowska, Robert Oostenveld, Anil M. Tuladhar, Britta U. Westner, Vitória Piai

## Abstract

White matter (WM) microstructure has been mainly associated with accuracy of picture naming. However, little is known about how WM properties relate to the *timing* of picture naming and its underlying electrophysiological activity. Here, we linked diffusion-weighted, electrophysiological, and behavioural data, using time-based measures to conceptually connect WM deterioration to alterations in conduction speed and precision. Our sample included individuals of varying age, with or without stroke. We correlated fractional anisotropy (FA) of bilateral WM tracts traversing the temporal lobe with mean response times (RTs) and response time variability (coefficient of variation, CV) in picture naming, and with the onset latency of neuronal activity, represented by the electrophysiological P2 component time-locked to picture onset. We selected tracts showing associations with P2 latency and naming performance separately, on which we conducted path analyses examining the relationship between structure, neuronal activity and behaviour. Depending on the tract, higher FA was associated with faster and/or more consistent naming, suggesting that ventral tracts differentially contribute to naming speed and consistency. Importantly, path modelling for the right inferior fronto-occipital fasciculus (IFOF) revealed that FA predicted P2 latency, which in turn predicted RTs, possibly reflecting a structure-activity-behaviour relationship in lexical-semantic processing. P2 source reconstruction demonstrated anatomical proximity to posterior segments of the right IFOF, supporting these findings. Overall, this study provides a proof of concept for modelling structure-activity-behaviour relationships in language research, while highlighting the need for elaboration of such relations at a more mechanistic level.

## 1. Introduction

The production of words is supported by a network of grey matter (GM) areas in the temporal, parietal and frontal cortices, which are connected via white matter (WM) association tracts (Dick et al., 2014; Shekari & Nozari, 2023). Given the role of WM in supporting rapid and precise neural communication across GM regions subserving cognition (Filley & Fields, 2016), a link between WM and the timing of word retrieval is expected. To date, however, relations between WM and word production have been predominantly investigated using accuracy measures (e.g., Houston et al., 2019; Ivanova et al., 2016). Moreover, when response time (RT) measures are reported in language WM research, they typically reflect mean naming times (e.g., Gold et al., 2007) but not naming time variability, another relevant measure of performance efficiency (e.g., MacDonald et al., 2003, 2006).

It is commonly assumed that neuronal activity, which can be measured using electrophysiology, travels along WM tracts from one GM area to another, ultimately resulting in behaviour. To date, however, few studies have examined the entire path of *WM structure – neuronal activity – observed behaviour*, with research mainly focusing on either the WM-behaviour or WM-neuronal activity relations. The present study aimed to explore the structure-activity-behaviour path for selected language-relevant association tracts, and to relate WM properties to picture naming performance.

### 1.1 WM, Neuronal Activity and Behaviour

White matter structure can be non-invasively characterised via diffusion MRI-derived measures such as fractional anisotropy (FA). In the adult brain, healthy WM is associated with higher FA, suggesting less diffusion and, hence, more organised and coherent myelinated fibre structures. FA decreases, by contrast, mark age-related changes (Beck et al., 2021; Lebel et al., 2012; Madden et al., 2012; Mendez Colmenares et al., 2023) as well as pathology (for review see Chanraud et al., 2010; Pinter et al., 2020), with cognitive function being impacted both in terms of accuracy and speed of performance (Coelho et al., 2021; Madden et al., 2012; Peter et al., 2025; Schellekens et al., 2025; Turken et al., 2008). In the language domain (e.g., Peter et al., 2025), FA in multiple, predominantly left-hemispheric association tracts has been linked to language production and comprehension performance (Houston et al., 2019; Ivanova et al., 2016; Troutman et al., 2022; for review, see Shekari & Nozari, 2023; Zhang et al., 2021).

Although the precise relationship between the physiological state of WM and microstructural proxies remains debated (e.g., Zhang et al., 2012), frequently suggested mechanisms involve myelination (Klawiter et al., 2011; Peters et al., 2019; but see Arshad et al., 2016). Given the role of myelin in electrical impulse conduction as well as regulation of conduction timing (Chorghay et al., 2018; Filley & Fields, 2016; Seidl, 2014), myelin changes likely affect communication between neurons by modulating the speed and temporal precision of the propagation of action potentials (e.g., Kim et al., 2013).

Signal conduction efficiency, (at least partly) modulated by the microstructural properties of WM, is expected to have an impact on the time course of neuronal activity. Previously, WM properties have been linked to temporal features of electrophysiological activity measured with electro-or magnetoencephalography (EEG or MEG) (e.g., Stufflebeam et al., 2008). For example, Takemura and colleagues (2020) investigated whether individuals’ peak latency of the visually evoked response (N75 component) in the visual cortex could be predicted by microstructural properties of the optic radiation tract, carrying information to the primary visual cortex. They found that WM properties explained 22% of the variance in peak latency of the N75 component. Importantly, there was no such relationship between the peak latency of the visually evoked component and a non-visual corticospinal control tract.

Similar approaches relating WM and electrophysiological activity have been applied in other cognitive domains (e.g., Berman et al., 2016; Westlye et al., 2009). Using a semantic task where participants judged whether pairs of words were related, Shin and colleagues (2019) investigated whether microstructure of the bilateral inferior longitudinal fasciculus was related to latency and amplitude of the semantic N400 component as well as to mean RT and accuracy in behavioural performance. They found correlations between the N400 and the microstructure of the left (latency and amplitude) and right (latency) inferior longitudinal fasciculus. A control auditory processing component showed no correlation with the WM tract of interest. Furthermore, WM properties did not correlate with behavioural measures, potentially due to the small sample size and/or the ceiling effects in performance. Overall, structure-activity studies of language processing remain scarce: Only few studies (e.g., Shin et al., 2019) have attempted to relate time-sensitive features of the electrophysiological signal to microstructural properties of language-relevant association WM tracts, and, to the best of our knowledge, no study has done so for language production.

Apart from neuronal processing, signal conduction efficiency is expected to have an impact on the observed behaviour. Overall, suboptimal conduction timing and precision might be sufficient to affect cognitive function without requiring complete conduction failure from severe demyelination or axonal injury (Filley & Fields, 2016; Pajevic et al., 2014). Thus, disruptions in conduction timing and precision remain a plausible mechanism behind the WM-driven age-, disease- and trauma-related deterioration of function (MacDonald et al., 2009).

Hypothetically, neural noise (random or irregular fluctuations that are not part of the signal and interfere with it) could impact neural processing and, consequently, behaviour in multiple ways. In one scenario, noise could lead to overall processing slowing, manifesting as a systematic, global shift in neural signal conduction speed. This overall neural slowing would subsequently result in cognitive slowing, reflected in RT measures of central tendency such as the mean and median. For instance, individuals with multiple sclerosis, a disease whose primary characteristic is demyelination, show longer mean RTs on a range of tasks, with the extent of slowing linked to the severity of disease (Kail, 1997). Importantly, when cognitive processing speed is controlled for, performance on memory or executive function tasks in this patient group is similar to the control group (e.g., Genova et al., 2013). In healthy participants, the association between WM and mean response speed has been reported for language tasks such as lexical decision (Gold et al., 2007). Regarding word production, mean-based RT investigations are scarce, with prior studies mainly using accuracy (e.g., Ivanova et al., 2016) or a joint accuracy-mean RT index (e.g., Troutman et al., 2022).

In another scenario, neural noise could lead to processing inconsistency manifesting as increased intra-individual, trial-to-trial variability (Faisal et al., 2008; MacDonald et al., 2006, 2009). This inconsistency in RTs can be captured by measures reflecting the spread of the RT distribution such as the standard deviation (SD) or the coefficient of variation (CV, SD divided by the mean). Similarly to the overall response speed, increases in intra-individual RT variability are observed in ageing and brain disorders, and have been argued to provide information about central nervous system integrity complementarily to central tendency measures (e.g., MacDonald et al., 2003, 2006). For instance, higher trial-to-trial variability has been reported to precede and predict greater performance decline in older individuals on tasks of processing speed, executive function, memory, and language, independently of the mean RT (Jutten et al., 2024; Lövdén et al., 2007; MacDonald et al., 2003). Multiple studies have reported correlations between microstructural properties of WM and variability in cognitive performance, particularly for processing speed and executive function (Fjell et al., 2011; Mazerolle et al., 2013; McCormick et al., 2023). Memory and language studies reporting RT variability are relatively scarce (e.g., Mella et al., 2013) and, to the best of our knowledge, the relation between WM properties and naming time variability is yet to be investigated.

It should be noted that the two hypothetical scenarios of neural noise impacting RTs are not mutually exclusive. Although a speaker might give slow but consistent (scenario one) or fast but inconsistent (scenario two) responses, in practice the individual mean RT and RT variability frequently correlate, and both have been related to poorer WM microstructure across the lifespan (e.g., McCormick et al., 2023). Nevertheless, these measures can predict performance independently and thus seem to reflect (at least partly) distinct underlying states or processes. Some authors have even suggested that performance variability, compared to central tendency measures, might be more sensitive to relationships between WM microstructure and, for instance, processing speed (Fjell et al., 2011; Mella et al., 2013; but see Booth et al., 2019).

### 1.2 Present Study

Given that WM properties might (partly) determine the levels of neural noise in the brain, thus impacting neuronal activity and behaviour, here we investigated the relationship between microstructural properties of ventral association tracts traversing the temporal lobe, neuronal activity implicated in word retrieval and picture naming performance. To this end, we analysed diffusion-weighted, EEG, and RT data during picture naming from stroke-free individuals and from individuals two months after left- (LH) or right-hemispheric (RH) stroke with no cognitive impairment. We included participants of a wide age range (26 – 80) and varying brain health to increase variability in our sample in order to study relationships between variables. Both presence of stroke and age were included as covariates in all our analyses. Efficiency of neuronal processing and of naming were defined using time-based measures, with shorter timing suggesting faster and more precise conduction of information along the WM tracts (i.e., less neural noise). Although some noise has been suggested to facilitate detection of weak signals (Wiesenfeld & Moss, 1995), thus potentially supporting processing in the brain (see MacDonald et al., 2006), here we conceptualised neural noise as detrimental to information processing (Kail, 1997; MacDonald et al., 2006; McCormick et al., 2023; Voytek et al., 2015).

Word production has typically been linked to the left-lateralised dorsal WM pathways (e.g., arcuate fasciculus, mapping acoustic information to the motor output systems). However, the temporal lobe has been shown to be critical for lexical access in word retrieval (Piai & Eikelboom, 2023; Shekari & Nozari, 2023). Here, we selected four ventral tracts, namely the inferior longitudinal fasciculus (ILF), inferior fronto-occipital fasciculus (IFOF), uncinate fasciculus (UF) and middle longitudinal fasciculus (MdLF) (see *Supplement*, https://osf.io/duxw6, Table S1-1 for anatomical description). In addition, we included the motor corticospinal tract (CST) as a non-language control tract (but see Forkel et al., 2022). All tracts were reconstructed bilaterally. Since signal conduction depends on multiple structural properties of WM (myelination, axonal diameter, internodal distance) whose relative contributions might be difficult to disentangle (Waxman, 1980), we chose a more general measure of FA, likely reflecting a combination of microstructural WM properties, including myelination (e.g., Song et al., 2003; Zhang et al., 2012).

Assuming that detrimental WM changes impact signal conduction timing and precision of neuronal communication, we selected mean RT (general speed) and coefficient of variation (RT variability) as behavioural measures of picture naming efficiency. The two measures potentially reflect different consequences of neural noise - overall slowing and inconsistent processing, respectively.

To operationalise the efficiency of neuronal signal conduction and neuronal communication, we calculated picture-locked ERPs and extracted the *onset latency* of the second positive peak, P2. The P2 component in picture naming (approximately 180-400 ms after picture onset) has a clear peak and is not contaminated by motor activity associated with articulation. It has been linked to lexical selection after semantic analysis of the picture (Costa et al., 2009; Rabovsky et al., 2021; Strijkers et al., 2010; for review see Piai & Borges, 2023) and might also be modulated by visual features of the stimuli (Roos et al., 2024). Previous MEG source reconstructions of picture naming activity between 200 and 400 ms have indicated multiple bilateral sources in the occipital and parietal lobes (Levelt et al., 1998; Liljeström et al., 2009), although these studies have not explicitly indicated whether this activity relates to the (MEG-counterpart) P2 component. Thus, to increase the validity and interpretability of our findings, we additionally reconstructed the underlying sources of the observed P2 component.

In sum, we used time-based measures to conceptually link electrophysiological and behavioural measures to signal conduction speed and precision, which are affected by WM deterioration. The P2 and ventral tracts are linked functionally via the association with visual and lexical-semantic processing, and anatomically via the occipito-parietal distribution (apart from UF). We hypothesised that the P2 reflects neuronal activity that can be related both to the WM microstructure of the ventral tracts and to naming performance (see **Figure 1**). We expected suboptimal signal conduction (as inferred from WM microstructure) to induce delayed P2 onsets, at least on some trials, leading to an overall delayed P2 onset in the averaged ERP signal. Subsequently, we expected longer P2 onset latencies to be associated with longer and/or more variable naming times, similarly to findings of P2 amplitude correlating with picture-naming responses (e.g., Costa et al., 2009).

**Figure 1.**
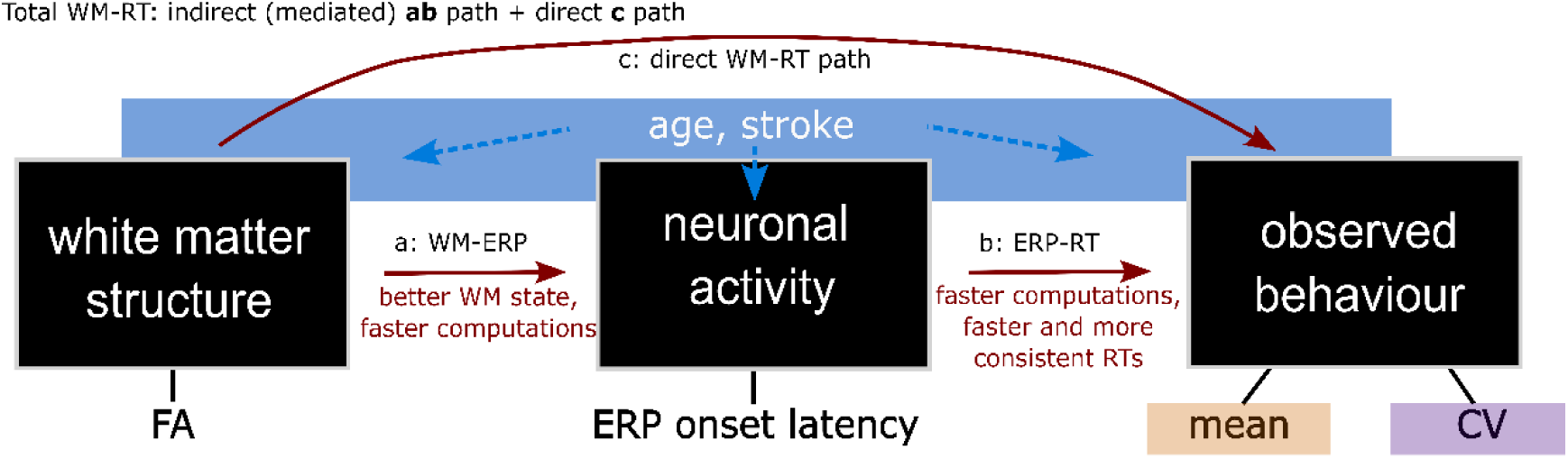
Structure-activity-behaviour (statistical) relationships assumed for path modelling. The black boxes are observation levels, with arrows indicating the path relationships (assumed to be causal) between them. **Path a**: The efficiency of neuronal computations (e.g., measured as ERP latency) reflects white matter (WM) properties. **Path b**: The efficiency of naming (RTs) is driven by the efficiency of computations. If **paths a and b** are both true, the neuronal activity bridges the structure-behaviour relationship (indirect **path ab**). In addition to this ERP-mediated path, WM might predict behaviour via variables other than the selected ERP measure (direct **path c**). The total impact of WM on behaviour is the combination of both paths (direct and indirect). Age and presence of stroke have been associated with measures at all three levels.

We first tested the structure-activity-behaviour relationships *successively* by running a series of partial correlations between WM, ERP, and behavioural measures. We performed the analyses for WM bilaterally, especially given the potential role of the right hemisphere in picture naming (e.g., Chupina et al., 2022; Levelt et al., 1998; Liljeström et al., 2009; Sörös et al., 2003). Next, based on the partial correlation results, we tested the structure-activity-behaviour relationships *simultaneously* by running path models for the tracts that showed correlations with both ERP and RT measures using structural equation modelling (SEM). We tested both the indirect path where WM and behaviour are linked via neuronal activity (path ab, **Figure 1**) as well as the direct path (c), which explains variance independent of P2 onset. For the ERPs to be considered an intermediary between WM structure and naming performance, the path model was expected to indicate the presence of relations in the indirect path (a: WM-ERP, b: ERP-behaviour).

## 2. Methods

The study was approved by the Medical ethics review committee (METC Regio Oost-Nederland, NL72600.091.20), following the Code of Ethics of the World Medical Association (Declaration of Helsinki). All participants received information in advance and gave written informed consent. Monetary compensation was provided for participation. The data were collected across two sessions in a longitudinal design at the Donders Centre for Cognitive Neuroimaging (DCCN, Radboud University) in Nijmegen, the Netherlands. Each session started with MRI scanning, followed by neuropsychological testing and experimental EEG. Here we report on the data from the first session only (for the original two-session study, see Roos et al., 2026).

### 2.1 Participants

Thirty-one right-handed native Dutch speakers (21 male, mean age 59, range 26 – 80 years) participated (see **Table 1**). Thirteen people with no history of stroke comprised the stroke-free group, which was matched in sex and education to the stroke group comprising 18 individuals (8 LH and 10 RH strokes). Participants for the stroke group were recruited at the department of Neurology, Radboud University Medical Center or the speech-and-language pathology department at the Canisius Wilhelmina hospital in Nijmegen and were diagnosed with an ischemic or haemorrhagic stroke of the middle cerebral artery based on CT or MRI by a certified neurologist. People with previous stroke history were included if the earlier stroke did not inflict long-term impairments.

**Table 1.** Demographic information for participants in the control and stroke groups (left-hemispheric, LH, and right-hemispheric, RH). Education is based on the 7-level classification of Verhage (1983), from unfinished primary school or special education (1) to higher vocational education (HBO) or bachelor’s degree (6), and university master’s degree or higher (7).

| group | sex | age:<br>mean, median, range | education level:<br>median, range |
| --- | --- | --- | --- |
| control | 9 m<br>4 f | 65, 70,<br>33 – 80 | 6,<br>3 – 7 |
| LH stroke | 5 m<br>3 f | 58, 65,<br>26 – 77 | 6,<br>2 – 7 |
| RH stroke | 7 m<br>3 f | 52, 54,<br>33 – 67 | 5,<br>4 – 7 |

*Note.* m = male, f = female.

The stroke lesion masks were manually delineated on T1- and T2-weighted images in ITK-SNAP (Yushkevich et al., 2006; www.itksnap.org) and normalised to MNI space, with the lesion overlap maps per hemisphere presented in **Figure 2**. Lesion volume (as obtained with MRIcroGL, v1.2.20220720, Rorden, 2025) was not statistically different (Welch independent samples t(8.4) = 0.53, *p* = 0.6) between the LH (N=7, mean = 14 cm^3^, SD = 12.3, range 1.3 – 34.8) and the RH (N=9, mean = 11.3 cm^3^, SD = 6.3, range 0.8 – 19) groups (lesions were not visible for one LH and one RH case, since the acute ischemic lesions were fully recovered during follow-up). Based on the Dutch version of the Boston Naming Test (Kaplan et al., 1983; Roomer et al., 2011), Amsterdam-Nijmegen Everyday Language Test (Ruiter et al., 2011) and the Token test (De Renzi & Faglioni, 1978), language deficits in the LH group were either absent or mild, with no participants meeting the cut-off for aphasia (score below 29 on Token test). Participants in the RH group did not show language deficits. All stroke participants were tested approximately two months post-stroke (mean 2.1 months, range 48 – 94 days).

**Figure 2.**
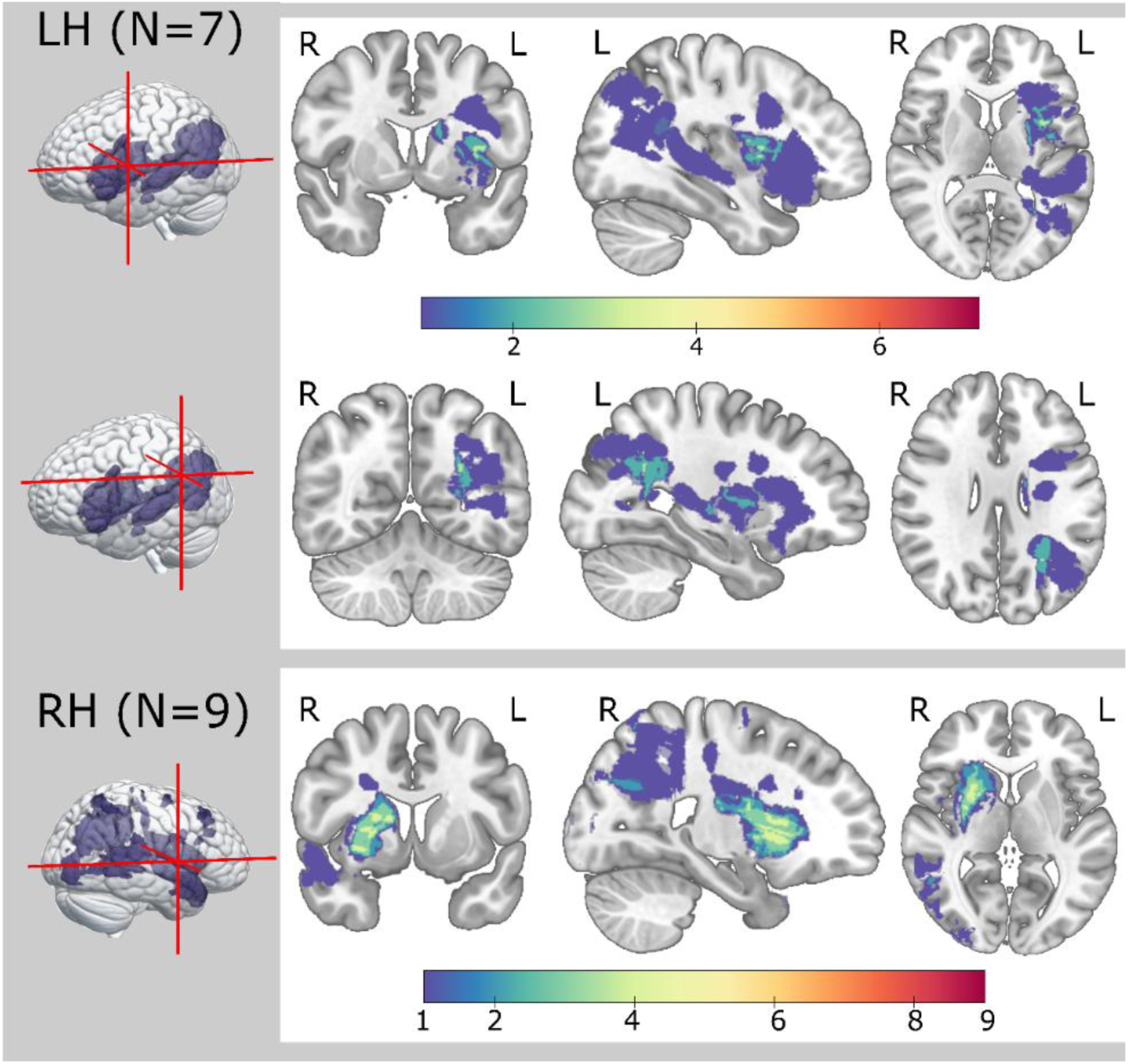
Lesion maps per stroke group. The colourbars indicate the number of participants. Lesion masks were drawn manually for participants with visible left-hemispheric (LH) and right-hemispheric (RH) lesions. Note that both groups included one participant where no lesion could be seen two months post-stroke (for more information, see Roos et al., 2026). The figure was created in MNI space using MRIcroGL (v1.2.20220720, Rorden, 2025).

### 2.2 Materials

The Concise Language Paradigm (CLaP, Roos et al., 2024) was used as the experimental EEG task. The CLaP consists of four experimental naming conditions (constrained, unconstrained, bare, scrambled), where pictures to be named are preceded by different types of auditory lead-in sentences (total N = 234). In the present study, we report the unconstrained picture naming condition for which the behavioural and electrophysiological responses were time-locked to and driven by picture presentation. This condition had more available trials (N = 78) while eliciting similar naming behaviour and electrophysiological responses as the bare picture naming condition (Chupina et al., 2022; Roos et al., 2024). In the unconstrained condition, participants heard pre-recorded neutral, semantically non-constraining sentences (cloze probability on the target 0-39%) read by a female native Dutch speaker. At the end of the sentence, a picture to be named was presented (e.g., *Het meisje opent het… [cadeau].* The girl opens the… [present]).

The pictures were coloured photographs of objects from the BOSS database (Brodeur et al., 2010) and internet. For more details, see Roos et al. (2024, 2026).

### 2.3 Design and Procedure

After the structural MRI scanning, EEG was recorded while participants performed the CLaP task. Participants were instructed to listen to the incomplete sentences and name the pictures presented at the end. Before the experiment, participants were familiarised with the stimuli pictures and completed several practice trials with different materials. Stimuli were presented in a unique list per participant using Mix (van Casteren & Davis, 2006). The experiment was presented with the Presentation software (Neurobehavioral Systems, Inc., www.neurobs.com). The full CLaP experiment comprised nine blocks of 26 trials each, with self-paced breaks after each block. Each trial started with a centred fixation cross on a black background that remained on the screen until picture presentation. After 1500 ms, the auditory lead-in sentence played (duration 1800 – 3600 ms, mean 2600 ms), followed by the 800-ms pre-picture interval. Next, the picture was presented for 2500 ms. The trial ended with a 2000-ms blinking interval marked by three asterisks. Responses were recorded from picture onset for 3500 ms using a microphone.

### 2.4 Structural Analysis

#### 2.4.1 MRI Acquisition and Preprocessing

T1, T2 and diffusion-weighted scans were acquired with a 32-channel head coil, either on a 3-T Siemens Magnetom Prisma or a PrismaFit scanner. After a localizer and automatic alignment scan to adjust the field of view, we acquired high-resolution anatomical T1 images with a magnetization prepared rapid acquisition gradient echo (MPRAGE) (TR = 2300 ms, TE = 3.03 ms, 192 slices of 1 mm, voxel size = 1 mm isotropic, FOV = 256 mm, acquisition time = 5m21s). Next, we acquired anatomical T2 fluid-attenuated inversion recovery (FLAIR) scans (TR = 5000 ms, TE = 397 ms, 192 slices of 1 mm, voxel size = 1 mm isotropic, FOV = 256 mm, acquisition time = 4m42s). Finally, we acquired a diffusion-weighted scan using a multiband echo planar imaging multi-shell sequence (TR = 3220 ms, TE = 74 ms, diffusion-encoding directions 30 × b = 1000 s/mm2 and 60 × b = 3000 s/mm2, 10 b = 0 images, multiband acceleration factor 3, acquisition time = 6m36s). We also acquired a b = 0 volume with inverted phase-encoding direction (acquisition time = 1m02s) for image distortion correction.

The T1- and T2-weighted DICOM images were converted to NIfTI format and bias-field corrected. For diffusion data preprocessing, skull-stripped images were used, with brain extraction performed using HD-BET (Isensee et al., 2019). Diffusion-weighted images underwent Marchenko-Pastur principal component analysis-based denoising and Gibbs artifact removal using MRtrix3 (version 3.0.0, www.mrtrix.org, Tournier et al., 2019). Next, susceptibility-induced distortions were corrected using TOPUP, and eddy current-induced distortions and head motion were corrected with EDDY (FMRIB Software Library, FSL, version 6.0.1, Jenkinson et al., 2012), including outlier replacement, slice-to-volume motion correction and susceptibility-by-movement interaction modelling (e.g., Dewenter et al., 2023).

#### 2.4.2 DWI Analysis

Tractography was performed in participants’ native space using FSL tools (FMRIB Software Library, FSL, version 6.0.1). First, we estimated fibre orientations in each voxel using BEDPOSTX (Behrens et al., 2007), which employs Markov Chain Monte Carlo sampling to model diffusion parameter distributions, particularly improving estimation in voxels with complex fibre configurations. Next, we performed probabilistic tractography with PROBTRACKX2 (Behrens et al., 2003, 2007) between seed and target regions of interest (ROIs) defining each tract. Exclusion masks were added to reject anatomically implausible pathways. The streamlines passing through both ROIs were considered to form the tract. Seed, target and exclusion masks for bilateral ILF, IFOF, UF and MdLF were obtained from XTRACT (Warrington et al., 2020). For ILF and MdLF, target and exclusion masks were adjusted in-house (for more detail, see van der Sluijs et al., in prep). Briefly, the target ROIs were positioned more posteriorly (middle of the parietal-occipital sulcus), with the MdLF mask additionally enlarged to include the inferior parts of the brain. Two new ROIs were added to the exclusion masks of each ILF and MdLF to avoid inclusion of the IFOF fibres. Bilateral CST ROIs were manually drawn on the template FA map from FSL, with seed in the motor cortex (precentral gyrus), target in the cerebral peduncle (midbrain) and exclusion being the brain midline separating the two hemispheres. To bring the masks from the standard to native space where tractography was conducted, transformation matrices between native (T1, diffusion) and standard MNI space (MNI152 T1 brain, 1 mm) were computed for each participant using FLIRT for linear (Jenkinson et al., 2002) and FNIRT for non-linear (Andersson et al., 2007) registration. PROBTRACKX was run bidirectionally between seed and target ROIs for each tract, with 10000 samples per direction (curvature threshold = 0.2, 2000 steps, step length = 0.5). Final tract probability density maps represent the sum of densities from both directions normalised by the total number of valid streamlines (waytotal). All tracts were thresholded at the 95^th^ percentile of streamline density (P95). The resulting tract masks were checked manually using *FSLeyes.* Artefactual single voxels were erased. The left MdLF did not yield a reliable reconstruction in one participant from the LH stroke group, most probably due to a significant disruption of the streamline paths by the lesion. Excluding this participant, the resulting sample comprised N=30 for all left MdLF analyses. Finally, we fit the diffusion tensor at the whole-brain level using DTIFIT to derive FA maps. By overlaying the probabilistic tract masks on the FA maps, measures of FA per tract and participant were extracted using *fslstats*. The derived FA measures were further used in the analyses as the estimates of the microstructural WM properties.

### 2.5 Neuronal Activity Analysis

#### 2.5.1 EEG Acquisition

EEG data were collected using Easycap electrode caps based on the equidistant M10 electrode layout (Easycap GmbH, Herrsching, Germany) with 64 actiCAP active electrodes (BrainProducts, Gilching, Germany). The data were recorded using BrainVision Recorder with an ActiChamp amplifier (500 Hz sampling rate, 0.1 – 200 Hz band-passed). We aimed at keeping the impedance at the ground (placed on the forehead) and reference (placed in standardised FCz position) electrodes below 5 kΩ and below 25 kΩ for the cap electrodes. To record electrooculograms (EOGs), we placed two external electrodes: one left of the left eye and one underneath the left eye. By combining the signal with the cap electrodes right of the right eye (corresponding to F10) and above the left eye (corresponding to Fp1), we computed horizontal and vertical EOGs during the offline analysis, respectively. Finally, we placed two additional electrodes above and below the right side of the mouth to record the electromyogram for detecting speech-related activity.

#### 2.5.2 EEG Analysis

##### 2.5.2.1 Preprocessing and ERP analysis

Preprocessing and analysis of the EEG data was performed in MATLAB (version R2021a) using FieldTrip (version 20231128, Oostenveld et al., 2011). Per participant, the continuous EEG data were segmented into trials from 800 ms before sentence onset (with sentences varying between 1800 – 3600 ms) to 400 ms after picture onset. All epochs were low-pass filtered at 50 Hz (twopass Butterworth IIR filter with order 6) and baseline-corrected using the 800 ms before sentence onset as the baseline window. Only trials marked as correct were analysed (see section 2.6). Based on visual inspection, we removed consistently noisy channels and extremely noisy trials. Next, independent component analysis was performed to detect eye-movement artifacts which were subsequently removed from the data. We further cleaned the data by removing noisy channels within single trials that exceeded a global rejection threshold: A channel was deleted from a trial if its signal SD exceeded the median SD across all channels and trials by more than two median absolute deviations. If more than half of the channels within the trial exceeded the threshold, the entire trial was discarded. Finally, all removed channels were interpolated based on the average of the neighbouring channels and inserted back into the data. After the data were re-referenced to the common average, the unconstrained naming trials were subset from the full CLaP dataset. The epochs were resegmented from –1000 to 400 ms relative to picture onset and low-pass filtered at 40 Hz. ERPs across trials per participant were calculated over 14 occipital electrodes (see **Figure 3**).

**Figure 3.**
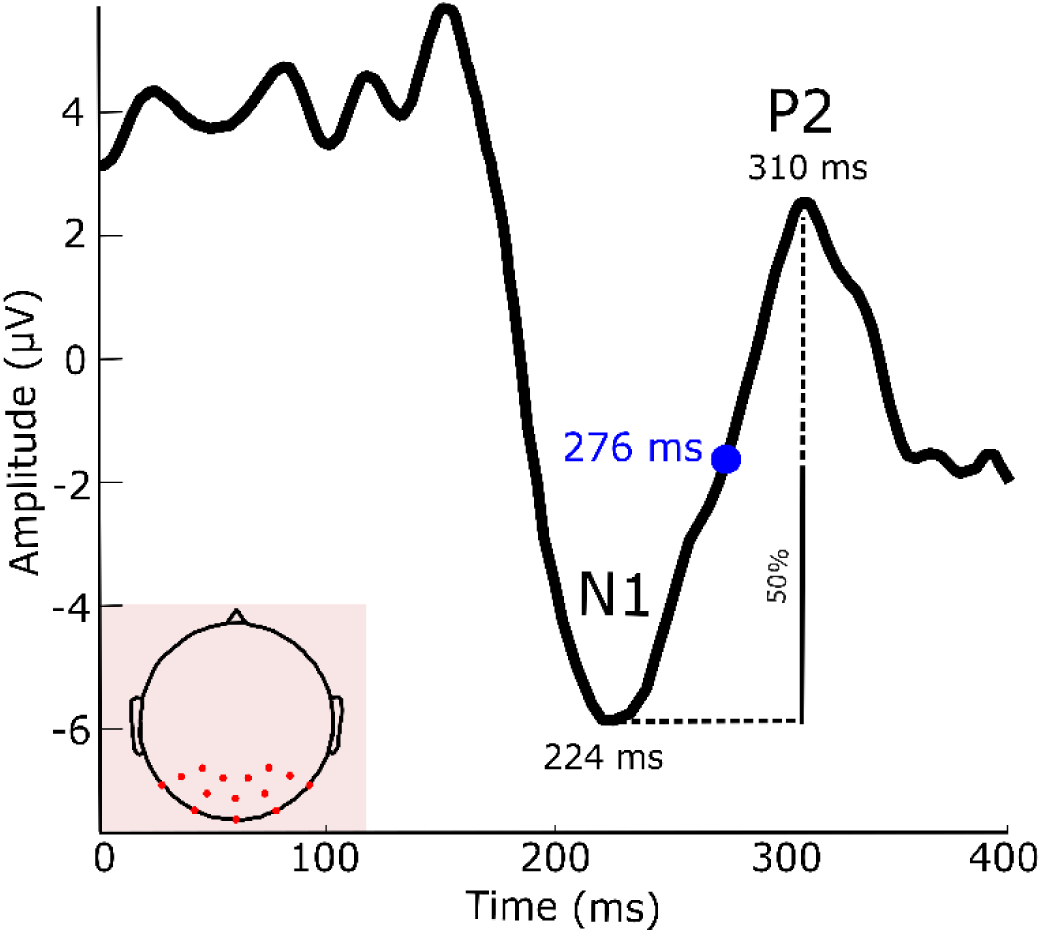
Example of an individual’s ERP waveform. ERPs were computed per participant over 14 occipital channels (see channel configuration in the left bottom corner). The half-peak P2 onset was calculated as the latency at the amplitude halfway between the preceding N1 trough and the P2 peak.

Peak amplitude and latency are two commonly used ERP measures. However, since ERPs are the product of trial averaging, the ERP peak amplitude can become smaller not only when the trial-to-trial amplitude is small, but also due to the latency of the peak varying from trial to trial. Even though it is possible to disentangle the true amplitude effects from the effects of the latency jitter (e.g., Walhovd et al., 2008), interpretation of the amplitudes across participants as a marker of neural or cognitive differences remains challenging, given that the amplitudes observed on scalp are highly affected by brain geometry and tissue conductivity. For instance, larger cavities filled with cerebrospinal fluid, typically observed in ageing and brain disease, can lead to attenuated or aberrant amplitudes that are not neurocognitively driven (Piai et al., 2024). Taking this into account, latency might be a more reliable measure than amplitude. Furthermore, onset latency of the component might be preferrable to peak latency, since the latter does not have a clear cognitive interpretation (Luck, 2014). Thus, we chose the P2 component onset quantified as half-peak onset latency as our ERP measure (Kiesel et al., 2008; Luck, 2014). In the ERPs of each participant, we first determined the minimum value within 140 – 250 ms after picture onset corresponding to the preceding N1 trough (see **Figure 3**). Next, we found a local peak (i.e., the maximum value) in the interval from the N1 trough to 400 ms marking the end of the ERP segment. Finally, we calculated the half-peak amplitude value between the N1 trough and P2 peak and extracted the corresponding latency. The derived half-peak P2 onset latency measures were further used in the analyses as a measure of neural processing efficiency.

##### 2.5.2.2 P2 source reconstruction

To determine the underlying cortical sources of the P2 ERP component in relation to the anatomy of the WM tracts, we performed source reconstruction using the finite element method (FEM) to take the stroke lesions into account. For this, we created detailed individual forward models using the FieldTrip SimBio software (Vorwerk et al., 2018). Forward models were based on high resolution tetrahedral meshes, derived from T1 and T2 structural MRI scans using simnibs (version 4.0.0) segmentation and meshing (*charm*, Puonti et al., 2020). Conductivity values were added per tissue type (WM = 0.22, GM = 0.47, csf = 1.71, scalp = 0.41, eye balls = 0.5, compact bone = 0.005, spongy bone = 0.048, blood = 0.57, muscle = 0.32, lesion = 0.75, where applicable; based on McCann et al., 2019), and dipoles were placed in the GM with approximately 3 mm resolution.

Electrode locations were recorded using a 3D scanner (Occipital Structure sensor, https://structure.io/, Homölle & Oostenveld, 2019) and co-registered to the MRI data. For more details on the source reconstruction methodology, please see Roos et al. (2026). We localised the ERP generators for the interval spanning between the half-peak onset latency (same as for the ERP latency measure) and the latency of the following P2 peak, determined per participant. This was contrasted with a pre-picture baseline window of the same length. We used a unit-noise-gain LCMV beamformer with a source orientation that maximises output power (van Drongelen et al., 1996; Van Veen et al., 1997) with common spatial filters for the P2 and the baseline segment. For the inversion of the covariance matrix in the beamformer computation, we used a truncated pseudo inverse and added 5% of sensor power to the diagonal of the matrix (Westner et al., 2022). Source-level power differences between both segments were evaluated using a Monte Carlo permutation approach (1,000 randomizations) based on dependent-samples t-tests (two-tailed) as implemented in FieldTrip.

### 2.6 Behavioural Analysis

Responses were coded for accuracy either online or offline using the recordings. RTs were measured manually using PRAAT (Boersma & Weenink, 2013). Target responses, synonyms (e.g., *kwast, penseel* for painting brush) and semantically related responses that adequately described the picture (*mond*, mouth instead of *lippen*, lips) were considered correct. All other responses were considered errors (omissions, silences, semantic or phonological errors, hesitations, anomias, multiple responses and responses that did not describe the picture adequately, e.g., *restaurant* instead of *train station*). The mean number of correct trials per participant was 75.8 (SD = 3.4, range 65 – 78). Unlike some previous studies (e.g., Fjell et al., 2011; Troutman et al., 2022), we did not remove the slowest and fastest responses to not decrease variability artificially.

Using correct responses, we calculated RT measures of central tendency and variability per participant. As a measure of central tendency reflecting overall naming efficiency, we used the *mean* RT. As a measure of naming variability, we used the *coefficient of variation* (CV), calculated by dividing the individual SD by the mean RT (e.g., Lövdén et al. 2007). The CV shows variability in RTs across trials while normalising it by the individual response speed, thus disentangling the two. The derived mean RT and CV measures were further used in the analyses as measures of behavioural performance efficiency.

### 2.7 Statistical Analyses

All descriptive and statistical analyses were conducted in R (v4.1.3; R Core Team, 2022).

#### 2.7.1 Structure-Activity-Behaviour Analyses

We ran a series of partial correlations between (i) WM (FA) and behavioural (RT mean, CV) measures, (ii) WM and ERP (P2 onset) measures, and (iii) ERP and behavioural measures. As there were no systematic demographic or performance differences between the LH and RH stroke groups (see **Table 1**, *Results* and *Supplement 2*, https://osf.io/duxw6), we combined them into a single stroke group for this and subsequent analyses. Thus, age and stroke status (control vs stroke group) entered into partial correlations as covariates. We ran both Spearman’s rank (*rho*) and Pearson’s (*r*) tests and checked for their convergence. While the non-parametric *rho* is our principal measure that ensures robustness by accounting for non-normally distributed variables and controlling for the impact of outliers, the parametric coefficient is frequently reported in the literature (e.g., Ivanova et al., 2016) and aligns with the linear modelling framework used in the path analyses. P-values for *rho* and *r* were adjusted for multiple comparisons using Holm’s correction (adjusted p-level < 0.05) for WM – RT tests (within central tendency and variability measures separately, across all 10 tracts) and for WM – ERP tests (across all 10 tracts).

Based on the correlation results, we tested the hypothesised structure-activity-behaviour relationship for right IFOF and left MdLF, the only two tracts for which FA measures simultaneously correlated with ERP latencies and RTs (as reported below). Using the SEM approach as implemented in the *lavaan* package (Rosseel, 2012), we fit three models (right IFOF mean, left MdLF mean and left MdLF CV) to estimate the parameters in a system of linear equations. In each model, we tested whether WM properties (FA) predicted ERPs (half-peak P2 onset latency, path a) and whether ERPs, in turn, predicted picture naming times (mean RT and CV, path b) (see **Figure 1**). In addition to this indirect mediation path (path ab), the models included a direct WM – RTs path (variance other than P2, path c) as well as the total WM – RTs effect (indirect and direct paths combined, ab+c). Age and group were included as covariates. All continuous variables were standardised and the categorical group variable was sum coded (–1 controls, 1 patients). All three models showed perfect fit (χ²(0), CFI = 1.00, TLI = 1.00, RMSEA = 0.00, SRMR = 0.00).

## 3. Results

Descriptive statistics for errors, RTs and P2 onset latencies are presented in **Table 2** (see also *Supplement 1*, Figure S1-1, https://osf.io/duxw6). FA measures per tract and group in relation to age are shown in **Figure 4** (for additional information, see *Supplement 2*, Table S2).

**Figure 4.**
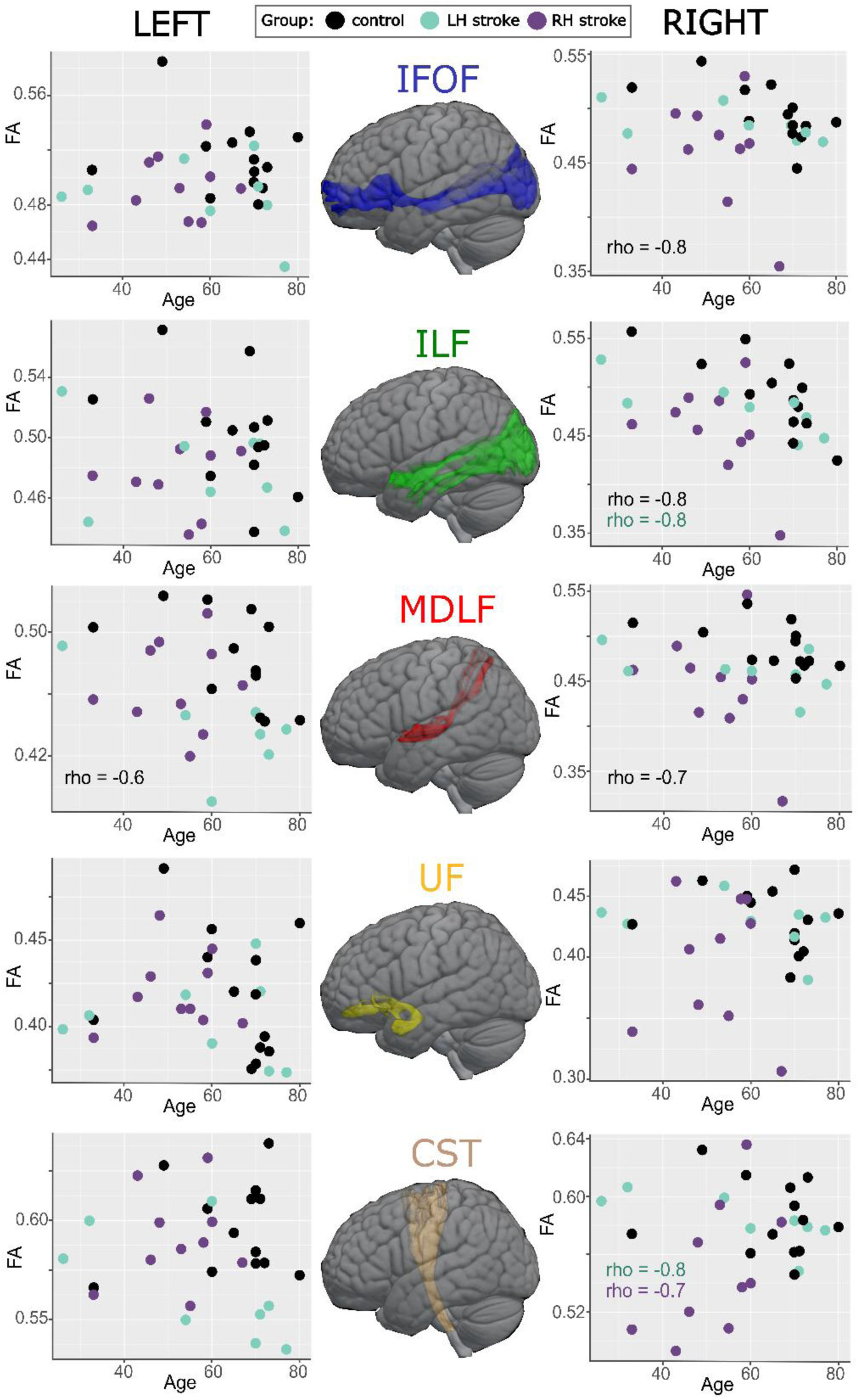
WM microstructure (FA) and age relationship. The scatterplots illustrate relations between FA and age per participant group (controls: black, LH stroke: mint, RH stroke: purple) for the inferior fronto-occipital fasciculus (IFOF), inferior longitudinal fasciculus (ILF), middle longitudinal fasciculus (MdLF), uncinate fasciculus (UF) and the control corticospinal tract (CST). The tract masks were obtained from the population-averaged tractography atlas by Yeh (2022) and plotted using MRIcroGL (v1.2.20220720, Rorden, 2025). The significant (alpha level < 0.05) non-parametric correlations between age and FA (tested within each group) are marked in the related tract subplots using the group colour (for more detail, see *Supplement 2*, Table S2).

**Table 2.** Naming error rates and RT measures (mean and coefficient of variation, CV), and P2 onset latencies per participant group.

|  |  | RT measures: group mean (SD),<br>range |  | ERP: group mean (SD),<br>range |
| --- | --- | --- | --- | --- |
| Group | Error rate, % | Mean (ms) | CV | P2 onset latency (ms) |
| Controls | 1.3 | 874 (93),<br>748 - 1080 | 0.26 (0.08),<br>0.13 – 0.42 | 245 (18)<br>218 – 276 |
| Stroke (all) | 4.2 | 1027 (127),<br>835 - 1298 | 0.3 (0.07),<br>0.19 – 0.39 | 240 (20)<br>199-284 |
| LH stroke | 5.3 | 1004 (112),<br>864 - 1158 | 0.32 (0.05),<br>0.27 – 0.38 | 239 (25)<br>199 – 274 |
| RH stroke | 3.3 | 1045 (141),<br>835 - 1298 | 0.29 (0.08),<br>0.19 – 0.39 | 241 (24)<br>206 – 284 |

### 3.1 Partial Correlations

After controlling for age and stroke status, FA in bilateral IFOF and right UF showed significant negative correlations with mean RT (corrected *p* < 0.005), FA in left CST with RT variability (CV, corrected *p* = 0.02), and FA in bilateral ILF and MdLF with both mean RT and variability (see **Table 3** and *Supplement*, Figure S1-2, https://osf.io/duxw6). The strength of all significant correlations was moderate (rho between –0.4 and –0.6). The non-parametric results converged with the Pearson’s test findings for all tracts but left ILF and IFOF (see *Supplement*, Table S1-2). ERP onset latency showed a significant correlation with mean RT (weak-moderate, rho = 0.34, *p* = 0.008) but not with RT variability (rho = 0.11, *p* = 0.39), converging with Pearson’s test results (see *Supplement*, Table S1-3). Finally, ERP onset latency was significantly negatively correlated with FA measures in the right IFOF, left UF and left MdLF (moderate, rho between –0.37 and –0.43, corrected *p*s < 0.03) (see **Table 4** and *Supplement*, Figure S1-2). Pearson’s test results converged for right IFOF and left UF but not left MdLF (see Supplement, Table S1-4).

**Table 3.** WM – RT partial correlations. Spearman’s rank (rho) correlations are reported with p-values adjusted across all tracts within central tendency and variability measures using Holm’s correction. The statistically significant (alpha < 0.05) correlations are marked in bold.

| tract | measure | rho | p-value |
| --- | --- | --- | --- |
| CST left | mean | –0.18 | 0.338 |
|  | CV | **–0.4** | **0.01** |
| CST right | mean | –0.27 | 0.098 |
|  | CV | –0.18 | 0.643 |
| ILF left | mean | **–0.44** | **0.003** |
|  | CV | **–0.46** | **0.002** |
| ILF right | mean | **–0.43** | **0.003** |
|  | CV | **–0.44** | **0.003** |
| IFOF left | mean | **–0.42** | **0.003** |
|  | CV | –0.17 | 0.643 |
| IFOF right | mean | **–0.53** | **<0.001** |
|  | CV | –0.27 | 0.18 |
| UF left | mean | –0.13 | 0.338 |
|  | CV | 0.17 | 0.643 |
| UF right | mean | **–0.41** | **0.005** |
|  | CV | 0.02 | 0.87 |
| MdLF left | mean | **–0.45** | **0.003** |
|  | CV | **–0.4** | **0.01** |
| MdLF right | mean | **–0.58** | **<0.001** |
|  | CV | **–0.51** | **<0.001** |

**Table 4.** WM – ERP partial correlations. Spearman’s rank (rho) correlations are reported with p-values adjusted for multiple comparisons across all tracts using Holm’s correction. Statistically significant findings are in bold.

| tract | rho | p-value |
| --- | --- | --- |
| CST left | –0.17 | 0.727 |
| CST right | –0.03 | 0.8 |
| ILF left | –0.16 | 0.727 |
| ILF right | –0.25 | 0.373 |
| IFOF left | –0.15 | 0.727 |
| IFOF right | **–0.43** | **0.006** |
| UF left | **–0.37** | **0.027** |
| UF right | –0.23 | 0.451 |
| MdLF left | **–0.38** | **0.027** |
| MDLF right | –0.23 | 0.451 |

### 3.2 Structure-Activity-Behaviour Path Modelling

Following the partial correlation findings, we selected the tracts whose FA measures correlated with ERP as well as RT measures and fit path models for them. This resulted in three models – the mean RT model for the right IFOF (N = 31) and the mean RT and RT variability models for the left MdLF (each N = 30).

All three path models showed that earlier P2 onset was predicted by younger age but not by stroke status (see also *Supplement*, Figure S1-1, https://osf.io/duxw6). In contrast, faster naming was predicted by the absence of stroke but not by age. Furthermore, earlier P2 onset predicted faster naming (**Table 5**).

**Table 5.** Structure-activity-behaviour regression estimates for right IFOF (mean RT) and left MdLF (mean RT, CV) models. Direct WM-RT relations are reported within FA – RT models. Total FA-RT relationship includes both direct and indirect (mediated via ERPs) pathways combined. Since FA-ERP results (P2 onset) for the left MdLF mean and CV models are identical, they are reported only in the mean RT model. The symbol ∼ means “predicted by”. Significant p-values are in bold.

| Predictor | Estimate | Std Error | z value | p-value |
| --- | --- | --- | --- | --- |
| right IFOF (mean) |  |  |  |  |
|  | P2 onset ~ |  |  |  |
| FA | −0.38 | 0.14 | −2.7 | **0.007** |
| Age | 0.46 | 0.12 | 3.8 | **<0.001** |
| Stroke status | −0.1 | 0.16 | −0.6 | 0.5 |
|  | R<sup>2</sup> = 0.43 |  |  |  |
|  | mean RT ~ |  |  |  |
| P2 onset | 0.39 | 0.18 | 2.2 | **0.03** |
| FA direct | −0.27 | 0.13 | −2.1 | **0.04** |
| Age | −0.07 | 0.15 | −0.47 | 0.64 |
| Stroke status | 0.5 | 0.15 | 3.3 | **0.001** |
|  | R<sup>2</sup> = 0.59 |  |  |  |
|  | FA ~ mean RT |  |  |  |
| indirect | −0.15 | 0.07 | −2.2 | **0.03** |
| total | −0.4 | 0.15 | −2.7 | **0.007** |
| left MdLF (mean) |  |  |  |  |
|  | P2 onset ~ |  |  |  |
| FA | −0.25 | 0.16 | −1.6 | 0.12 |
| Age | 0.43 | 0.15 | 2.8 | **0.005** |
| Stroke status | −0.06 | 0.19 | −0.3 | 0.75 |
|  | R<sup>2</sup> = 0.31 |  |  |  |
|  | mean RT ~ |  |  |  |
| P2 onset | 0.49 | 0.17 | 3 | **0.003** |
| FA direct | −0.03 | 0.15 | −0.18 | 0.86 |
| Age | −0.05 | 0.16 | −0.3 | 0.77 |
| Stroke status | 0.58 | 0.19 | 3.1 | **0.002** |
|  | R<sup>2</sup> = 0.55 |  |  |  |
|  | FA ~ mean RT |  |  |  |
| indirect | −0.12 | 0.08 | −1.55 | 0.12 |
| total | −0.15 | 0.18 | −0.82 | 0.4 |

| left MdLF (CV) |  |  |  |  |
| --- | --- | --- | --- | --- |
|  | CV RT ~ |  |  |  |
| *P2 onset* | -0.11 | 0.2 | -0.5 | 0.6 |
| *FA direct* | -0.25 | 0.2 | -1.2 | 0.2 |
| *Age* | 0.27 | 0.18 | 1.5 | 0.14 |
| *Stroke status* | 0.26 | 0.23 | 1.2 | 0.25 |
|  | R<sup>2</sup> = 0.22 |  |  |  |
|  | FA ~ CV RT |  |  |  |
| indirect | 0.03 | 0.06 | -0.45 | 0.65 |
| total | -0.23 | 0.2 | -1.1 | 0.26 |

Regarding the impact of WM, in the mean RT model for right IFOF, higher FA predicted earlier P2 onset (β = −0.38, p = 0.007), with the full FA-ERP model explaining 43% of the variance. Next, higher FA also predicted faster naming (direct path, β = −0.27, p = 0.04), with the full model explaining 59% of variance. The indirect FA-RT relationship mediated by P2 was also statistically significant (β = −0.15, p = 0.03), with a total effect of FA on mean naming times (β = −0.4, p = 0.007). These results indicate that microstructural properties of the right IFOF were related both to earlier P2 onset and faster mean picture naming, with some variable(s) other than ERP latency explaining part of the variance in the WM-RT relationship.

In both mean RT and RT variability models for the left MdLF, WM did not predict ERP onset or naming performance. While this result seems at odds with the initial Spearman correlation findings, it could be explained by the fact that the effects for the left MdLF were less robust, that is, left MdLF – ERP relations did not survive the Holm’s adjustments in Pearson’s tests and the path models rely on linear estimation (*Supplement*, Table S1-4 and Figure S1-2).

### 3.3 P2 Sources and Path Modelling Findings

Source-level power was significantly different between the P2 interval (from half-peak onset to peak latency) and the baseline segment (*p* = .002). Sources underlying the P2 component at the group level are shown as thresholded t-values in **Figure 5**, corresponding to the maximum ∼20% of the distribution. The most prominent sources (t ≤ −5.49, 99^th^ percentile) were located in right occipital lobe, right angular gyrus, right cuneus and right inferior parietal lobule, with additional bilateral sources in pre/postcentral regions, (pre-)cuneus, inferior and superior parietal regions. For a complete list of grid-points with their corresponding AAL atlas parcel (Tzourio-Mazoyer et al., 2002), see *Supplement*, Table S1-5 (https://osf.io/duxw6).

**Figure 5.**
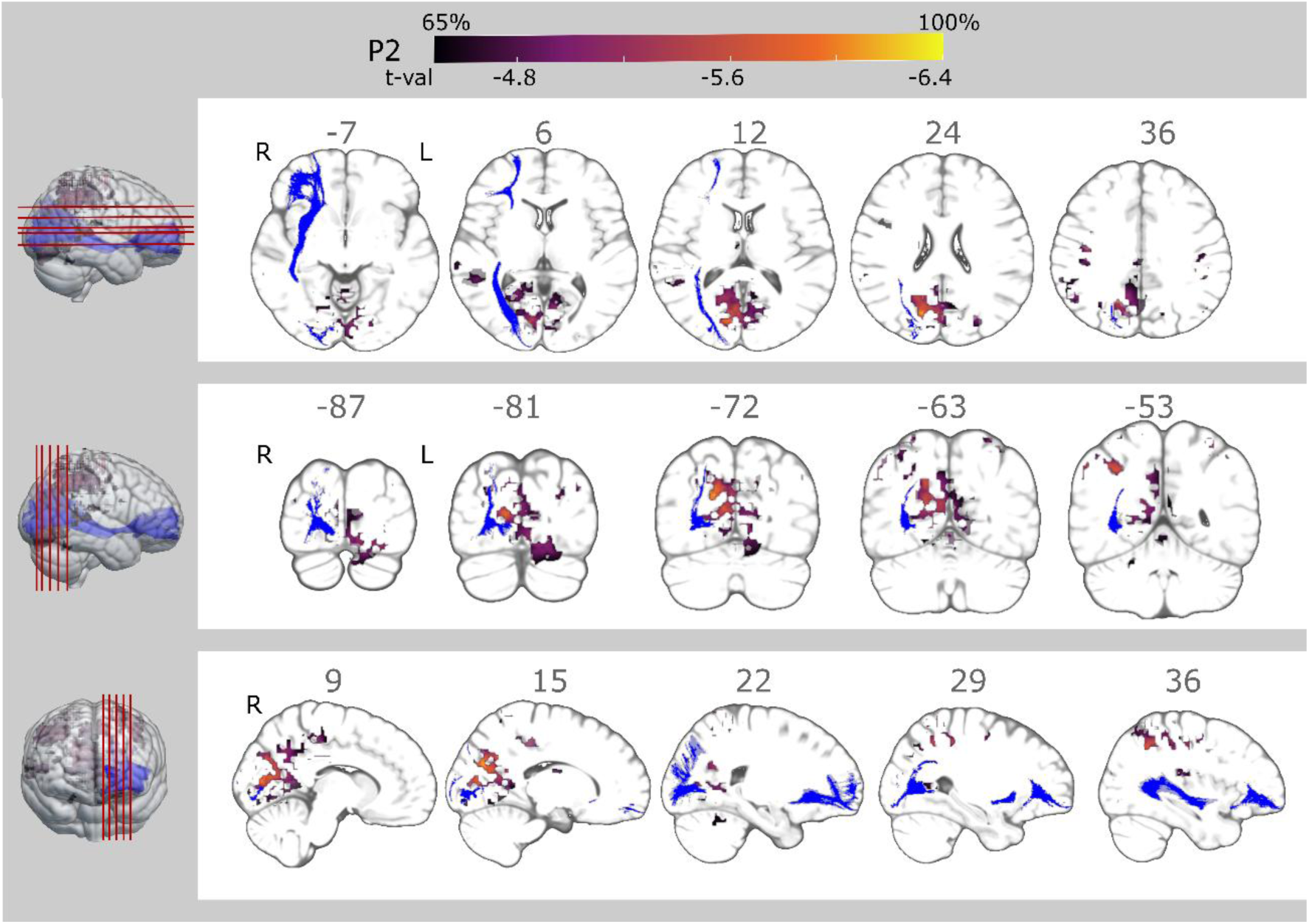
Location of the P2 sources (warm colours) and right IFOF (blue) at the group level are in close anatomical proximity. The slices with P2 sources and right IFOF are plotted on the axial (top panel), coronal (middle panel) and sagittal (bottom panel) planes. We thresholded the P2 sources at 65% of the maximum absolute t-value (t ≤ −4.1; minimum t-value = −6.4), corresponding to the upper ∼20% of the distribution. The IFOF mask is obtained from the population-averaged tractography atlas by Yeh (2022). The plot was created using MRIcroGL (v1.2.20220720, Rorden, 2025). For 3D representation of the P2 sources and their anatomical proximity to the right IFOF, see https://osf.io/duxw6.

The largely parieto-occipital distribution of the P2 sources with a right-hemisphere bias is well aligned with the path modelling results, suggesting an association between the P2 (onset latency) and WM microstructure of the right IFOF. The anatomical proximity between these P2 sources and the posterior portions of the right IFOF, illustrated in **Figure 5**, further corroborates our path modelling findings (for videos with a 3D representation of the P2 sources and their location in relation to the right IFOF, see https://osf.io/duxw6).

## 4. Discussion

We investigated whether microstructural properties of the bilateral ventral WM tracts predict both picture naming performance and associated electrophysiological activity using a sample of stroke-free and post-stroke individuals. By using time-based behavioural (mean RT, CV) and electrophysiological (ERP onset) measures, we conceptually connected these measures to the consequences of WM deterioration, that is, alterations in conduction speed and precision (e.g., Filley & Fields, 2016). We found that while most of the bilateral ventral tracts were linked to picture naming performance, their microstructure differentially related to naming speed and/or variability. Importantly, we found a full WM structure-neuronal activity-behaviour relationship linking the microstructural properties of the right IFOF and the speed of picture naming performance directly *and* indirectly via the parieto-occipital electrophysiological P2 component. Finally, we showed that the later onset of the P2 component was linked to both advanced age and slower picture naming. Below, we discuss these findings in more detail.

### 4.1 Structure-Activity-Behaviour Relationship: Microstructure of Right IFOF Predicts Picture Naming Speed via P2

We found a structure-activity-behaviour relationship such that higher FA of the right IFOF predicted earlier P2 onset which, in turn, predicted faster picture naming. The anatomical proximity between the electrophysiological P2 sources and the posterior portions of the right IFOF in occipito-parietal areas lends further credibility to this neuro-behavioural pathway (**Figure 5** and https://osf.io/duxw6). Due to associations of each of the three levels with lexical-semantic processing, this pathway likely underlies lexical access during word planning, as we discuss next.

The prominent role of the IFOF in language production has been linked to its involvement in visual and lexical-semantic processing (Cocquyt et al., 2020; Shekari & Nozari, 2023). Compared to other ventral tracts, the left IFOF shows the strongest and most consistent correlations with post-stroke naming accuracy (Zhang et al., 2021). Anatomically, the IFOF extends from posterior to anterior areas, connecting occipital, parietal and temporal cortices with frontal areas, including the inferior frontal gyrus (Radwan et al., 2022; Shekari & Nozari, 2023). These are the regions that are also associated with brain activity during picture naming, proceeding from visual processing in posterior areas towards phonological and articulatory processes in frontal areas (de Zubicaray, 2023; Indefrey, 2011; Roelofs, 2014). The cortical sources of the electrophysiological P2 component in our study were localised in particular to the occipito-parietal cortex, in line with previous MEG findings (Levelt et al., 1998; Liljeström et al., 2009; Sörös et al., 2003). Analogously to the IFOF, the P2 has been linked to lexical selection (Costa et al., 2009; Strijkers et al., 2010; for review see Piai & Borges, 2023) and seems to be modulated by the visual and name agreement properties of the picture stimuli (Cheng et al., 2010; Roos et al., 2024). Finally, picture naming speed is determined by the accumulation of processing times of all stages, starting from visual picture onset in occipital areas up until articulation in frontal and pericentral areas, thereby capturing all processes required for naming. Altogether, these are potentially the combined reason why we observed this structure-activity-behaviour relationship with the IFOF, rather than with other tracts.

In addition to the indirect connection of WM and naming performance via the P2, part of the variance between WM and naming times was explained by the direct WM-RT path, indicating impact of other variables not tested in the model, such as GM properties and/or processes linked to other stages of word retrieval. This is to be expected, as it would not be very plausible if P2 latency explained all of the variance in the WM-RT relationship. For the left MdLF, the structure-activity-behaviour relationship was neither found in the mean nor in the variability model. Apart from the explanation that linear path models were not suitable to capture this relationship for the left MdLF, this result might also stem from the tract being less functionally relevant for lexical access during word retrieval (e.g., Shekari & Nozari, 2023).

While both left and right IFOF correlated with mean naming times, a relation with the P2 component was only observed for the right IFOF. This might seem surprising given the critical role of LH neural structures for language performance (e.g., Ivanova et al., 2016; Ojemann, 1983; Wilson & Schneck, 2020; Zhang et al., 2021). An involvement of RH tracts is however consistent with the contribution of the RH to language in both healthy and clinical populations (e.g., Chang & Lambon Ralph, 2020), specifically supporting the notion that the ventral cortical areas and tracts subserve (lexical-semantic) processing bilaterally (e.g., Cocquyt et al., 2020; Hickok & Poeppel, 2007). The lack of observing a structure-activity-behaviour relationship for the LH might thus be due to our sample. We included LH stroke cases tested in the subacute phase, which might have introduced a bias towards the RH (Saur et al., 2006). Thus, while involvement of the IFOF (and other ventral WM tracts) in picture naming seems to be certain, it remains to be investigated whether the entire structure-activity-behaviour relationship observed for the right IFOF can be replicated in the LH in a sample without LH strokes. Currently, our results suggest that the RH seems to support language computations in a way that can impact the timing of word production, especially when the LH is lesioned.

### 4.2 Mean Naming Speed and Variability are Differentially Associated with Ventral WM-Pathways

We found moderate correlations between FA in most bilateral association tracts and RT measures, with WM tracts differentially related to either mean RT, CV, or both measures. Bilateral IFOF and right UF were related to mean naming times and bilateral ILF and MdLF to both mean naming times and naming time variability. The only tract purely associated with RT variability was left CST. The consistency of these patterns across the hemispheres for ILF, IFOF, and MdLF suggests a degree of (functional) similarity for the homologous tracts, in line with previous findings that homologous ventral tracts exhibit high correlations across microstructural measures (Wahl et al., 2010). Overall, our results are consistent with previous findings largely based on accuracy (for review see Shekari & Nozari, 2023; Zhang et al., 2021). Since response speed in picture naming is strongly linked to efficiency of lexical access (e.g., Costa et al., 2009; Laganaro et al., 2012), the associations between bilateral ILF, IFOF, and MdLF and mean RT further support their functional relevance to this process.

Interpretation of differential association of the tracts with mean RT or RT variability remains challenging, since the mechanisms behind these measures are not clearly established. Following Jensen (1992), we assume that speed of processing (mean RT) reflects the average neural conduction velocity in the WM association pathways, with general performance slowing resulting from systematic noise along the word retrieval path. The level of this noise and its disruptiveness to processing, however, are likely determined by multiple variables and not exclusively the WM properties. For RT variability, we speculate that it is driven by the combination of more sporadic neuronal noise along the WM paths (i.e., random errors generated while neurons conduct the signal) and/or the oscillatory states of the neural system indicating readiness to process the stimulus (e.g., Faisal et al., 2008; Jensen, 1992; Paraskevopoulou et al., 2021). Interestingly, Jensen (1992) also speculated that axonal properties (e.g., diameter) and myelination could underlie the two RT measures: Both are associated with signal conduction speed enabling faster processing, while myelination further decreases neural noise by controlling “leakage” of action potentials along the WM. Like the correlated central tendency and variability measures, axonal and myelin properties are interdependent but also distinct, and can therefore potentially explain differential results for mean RT and RT variability.

### 4.3 Later P2 Onset is Predicted by Older Age and Predicts Slower Naming

Our path models showed no systematic impact of stroke status on P2 onset latency. This might be attributed to the instantaneous nature of volume conduction, the heterogeneous lesion locations and the fact that the most reliable P2 sources were found in occipito-parietal areas, which were intact in most participants (**Figures 2 and 6**, also see Piai et al., 2024). That is, the scalp-level P2 onset latency estimates are unlikely to be impacted by the stroke lesions in our sample. By contrast, later onset of the P2 component in picture naming was associated with advanced age. In prior literature, older age has been linked to ERP amplitude changes and increased latency, which are linked to lower efficiency of behavioural performance (e.g., Boutet et al., 2021; Cespón et al., 2013; Gajewski et al., 2018). Some language studies have reported delayed peak latency in older age for early sensory (e.g., Federmeier et al., 2003) or both sensory and language components (for review, see Wlotko et al., 2010). Our study provides further evidence that language-related electrophysiological components, namely the component associated with lexical processing in picture naming, might show later onsets in older age. Importantly, similarly to the amplitude modulations in the P2 time window (200 – 400 ms) (e.g., Costa et al., 2009; Laganaro et al., 2012), later P2 onset predicted slower picture naming responses in the current study, thus connecting time-resolved features of the neuronal signal to the timing of behavioural performance.

### 4.4 Limitations and Future Directions

In the present study, we hypothesised that microstructural properties of the ventral WM tracts would impact signal propagation across brain areas, which, in turn, would impact the electrophysiological signatures of neuronal processing. In line with this assumption, Park et al. (2024) demonstrated that signals seem to propagate less efficiently from visual cortex to other regions when microstructural properties of the connecting WM were less optimal across the participant sample. While these simplified models seem to work for empirical investigations, they are unlikely to do justice to the complexity of signal propagation and neuronal activity in the brain. For instance, electrophysiological activity along the ventral stream can exhibit cortico-cortical signal propagation (Woolnough et al., 2022) as well as recurrent and feedback propagation (Eickhoff et al., 2025; Riesenhuber & Poggio, 2000). The latency of the P2 likely depends on properties of local and connected GM in areas where the information is processed. Furthermore, since microstructural WM properties might change along the tract (e.g., Takemura et al., 2020), associations can differ per section of the tract. Such factors could explain additional variance not captured with our simplified model. Taking these considerations into account, future investigations would benefit from developing more detailed structure-activity-behaviour models by combining multiple anatomically and electrophysiologically relevant variables.

Next, we used DTI-derived FA as a proxy for microstructural properties of WM. FA is derived by combining the three eigenvalues of the diffusion tensor and, thus, can be considered a general measure of WM. While the determinants of microstructural WM properties remain debated, FA has been suggested to mainly reflect either axonal or a combination of axonal and myelin integrity (Peters et al., 2019; Song et al., 2003; Zhang et al., 2012). Other measures of WM microstructure such as axial, radial, and mean diffusivity could be used together with FA to provide complementary information and might be able to capture aspects of axonal health or myelination more specifically, compared to the more general FA index (e.g., Jones et al., 2013; Peters et al., 2019; Song et al., 2003; Zhang et al., 2012). It is possible that more myelin-sensitive measures such as radial diffusivity might reveal stronger structure-activity-behaviour relationships, given that propagation of action potentials is highly dependent on myelin (e.g., Kim et al., 2013). Our findings could be taken as the basis for more hypothesis-driven, confirmatory studies using these additional measures.

Finally, we found that P2 onset predicted mean naming times but not naming time variability. This is not surprising given that individual half-peak onset measures were derived from the ERP, an averaged waveform, likely reflecting the central tendency measures of the P2 component across trials. To obtain a variability-based measure of the component onset to connect structure and neuronal activity to naming time variability, future studies could attempt to determine the onset of the P2 (given sufficient signal-to-noise ratio) at the trial level and calculate the averaged onset latency based on those values.

## 5. Conclusion

We aimed to relate the microstructure of language-relevant ventral WM tracts to time-based measures of neuronal activity and picture naming performance. Mean naming times correlated with WM properties of the bilateral ILF, IFOF, MdLF, and right UF, but not left UF or the control motor CST. Bilateral ILF and MdLF were also linked to low variability of response times, thus demonstrating that naming time speed and variability can dissociate with respect to the associated structural connectivity. While the neural mechanisms for mean naming speed and variability remain to be elucidated, mean RT can be speculatively connected to the overall properties of WM and GM, whereas RT variability may be connected to the degree of sporadic noise and/or readiness of the neural system to process the propagating signal. Regarding the full structure-activity-behaviour path, we found that P2 onset latency was predicted by WM properties of the right IFOF, revealing a significant relationship between structural connectivity, neuronal activity, and mean picture naming times for this tract. In fact, all three observation levels are functionally connected via their contribution to lexical-semantic processing (in terms of its timing). In addition, reconstruction of the P2 to occipito-parietal sources revealed close anatomical proximity to the posterior parts of the IFOF, lending more credibility to these results. While our findings serve as a proof of concept for investigating how structural connectivity, neuronal activity and behavioural performance are related, future research is required to establish the connection between WM, underlying electrophysiological computations and behaviour resulting from them.

The **supplementary materials** can be found at https://osf.io/duxw6.

## Acknowledgements

We are very grateful to **Anna Dewenter** for help with setting up the pipeline for diffusion-weighted data preprocessing. We also would like to thank all the Master’s students from Radboud University who contributed to the longitudinal data collection effort.

## Author contributions (CRediT)

**Irina Chupina**: Conceptualisation; methodology; formal analysis; writing – original draft; writing – review and editing; visualisation. **Natascha M. Roos**: Conceptualisation; methodology; formal analysis; writing – review and editing; visualisation; data curation; investigation. **Joanna Sierpowska**: Methodology; writing – review and editing. **Robert Oostenveld**: Conceptualisation; methodology; supervision; writing – review and editing. **Anil M. Tuladhar**: Resources; writing – review and editing. **Britta U. Westner**: Conceptualisation; methodology; supervision; writing – review and editing. **Vitória Piai**: Conceptualisation; methodology; writing – review and editing; supervision; project administration; funding acquisition.

